# Determinants of birth asphyxia among newborns in Amhara national regional state referral hospitals, Ethiopia

**DOI:** 10.1101/649467

**Authors:** Alemwork Desta Meshesha, Muluken Azage, Endalkachew Worku, Getahun Gebre Bogale

**Author notes:** Corresponding author (AD).

## Abstract

**Background:** Globally, every year, 2.5 million infants die within their first month of life. Neonatal asphyxia is the leading specific cause of neonatal mortality in low- and middle-income countries, including Ethiopia. Therefore, the aim of this study was to identify the determinants of birth asphyxia among newborns admitted in Amhara region referral hospitals, Ethiopia.

**Methods:** Facility-based unmatched case-control study was employed among 193 cases and 193 controls of newborns. Newborns admitted to neonatal intensive care units with admission criteria of birth asphyxia and without birth asphyxia were considered as cases (Apgar score<7) and controls (Apgar score>=7) respectively. Data were collected using a structured questionnaire by systematic random sampling technique with proportional allocation, and entered in to Epi-Info version 7 and exported to SPSS version 20 for statistical analysis. Bivariate and multivariable logistic regression models were fitted to identify determinants of birth asphyxia.

**Results:** Newborns with low birth weight (<2.5kg) had 8.94 higher odds of birth asphyxia than those whose weight at birth was >=2.5kg at birth (AOR: 8.94, 95% CI: 4.08, 19.56). Newborns born at health centers were 7.36 times more likely to develop birth asphyxia than those born at hospitals (AOR: 7.36, 95% CI: 2.44, 22.13). Newborns born using instrumental delivery were 3.03 times more likely to develop birth asphyxia than those delivered by vaginally (AOR: 3.03, 95% CI: 1.41, 6.49). Newborns from mothers with prolonged labor were 2 times more likely to suffer from birth asphyxia as compared to their counterparts (AOR: 2.00, 95% CI: 1.20, 3.36).

**Conclusion:** This study identified prolonged labor, instrumental delivery, delivered at health centers, and low birth weight were identified as determinants of birth asphyxia. Thus, intervention planning towards the identified determinants may be needed to reduce neonatal birth asphyxia.

## Introduction

Birth asphyxia is a condition characterized by an impairment of exchange of the respiratory gases (oxygen and carbon dioxide) resulting in hypoxemia and hypercapnia, accompanied by metabolic acidosis [1]. Birth asphyxia is defined by the World Health Organization as “the failure to initiate and sustain breathing at birth”. Worldwide, birth asphyxia is a serious clinical problem leading to significant mortality and morbidity. Each year approximately 24% of neonatal deaths occurred due to birth asphyxia with an equal number of survivors with serious neurological sequelaes, such as cerebral palsy, mental retardation and epilepsy leading to detrimental long term consequences for both child and family [2]. Birth asphyxia leads to the impairment of normal exchange of respiratory gases during the birth process and subsequent adverse effects on fetus [3]. International reports indicated that birth asphyxia is the third cause of neonatal deaths (23%) next to infections (36%) and preterm (28%) [4]. In Ethiopia, birth asphyxia contributed 24% of neonatal deaths [5]. The Amhara national regional state health bureau estimated for 2017/18 that its prevalence was 7.8%.

Mothers and their newborns are vulnerable to illnesses and deaths during the postnatal period [6]. Worldwide, 2.5 million infants die within their first month of life every year, contributing nearly 47% of all deaths of children under-five year’s age. Almost all deaths of newborns are in developing countries, with the highest number in South Asia and sub-Saharan Africa [7]. Birth asphyxia is the leading specific cause of neonatal mortality in low and middle-income countries and it is also the main cause of long-term illnesses including mental retardation, cerebral palsy, and other neurodevelopmental disorders [8]. Ethiopia is one of the ten countries with the highest number of neonatal mortality worldwide, with an estimated number of 122,000 newborn deaths per year [9].

Studies from abroad indicated that low birth weight, caesarian section [10-12]; multiple births, lack of antenatal care[13]; maternal age, gravidity, mode of delivery [14], and prolonged labor, meconium stained amniotic fluid and fetal distress [12] were the significant causes of birth asphyxia. However, most of the studies were restricted in single institution and based on secondary data or records which may face to data/info incompleteness. Therefore, the aim of this study was to identify significant determinants of birth asphyxia among newborns to formulate intervention mechanisms at local, regional and national level.

## Methods

### Study design, period and setting

Facility-based unmatched case-control study was employed to identify the determinants of birth asphyxia among newborns in Amhara national regional state referral hospitals from March 1 to April 30, 2018.

The Amhara National Regional State is located in the North Western part of Ethiopia between 9°20’ and 14°20’ North latitude and 36° 20’ and 40° 20’ East longitude. The Central Statistics Agency’s total population projection estimate for the Amhara Region for 2017 is 21,134,988 with a fifty-fifty numerical split between the sexes. Of these 17% were urban residents which are below the national average [15]. According to Ethiopian 2009 Ethiopian Fiscal Year (EFY) Annual Performance Report published by Federal Ministry of Health, Amhara has 68 Hospitals, 841 Health Centers and 3,342 Health Posts [16]. Among the sixty eight functional hospitals in the region, Dessie, Felege-Hiwot, University of Gondar, Debebirhan, and Debremaros hospitals are tertiary care (referral) hospitals. Thus, all the five referral hospitals are serving for all population found in the region.

### Study participants

All asphyxiated and non-asphyxiated newborns admitted to neonatal intensive care units (NICU) of Referral Hospitals found in Amhara Regional state are study population. Newborns diagnosed as birth asphyxia from NICU were included for cases, while newborns without birth asphyxia were counted for controls. Newborns with no mothers (caregivers) due to death or newborns with loss of mothers and mothers who are sick and unable to respond were excluded from the study.

### Sample size and sampling procedures

Sample size was calculated based on unmatched case control formula (Kelsey) with the assumptions of power=80% and 95% CI using Epi Info version 7. From previous case control studies on determinants of birth asphyxia, the major determinants were low birth weight (*p*=11.33%, *OR*=2.40), gestational age of <37weeks (*p*=52%, *OR*=2.57), multiple births (*p*=6.2%, *OR*=0.11), mode of delivery (*p*=22.2%, *OR*=2.94), and gravidity (*p*=33.3%, *OR*=2.64) [10, 14, 17]. From the alternative sample sizes, the largest sample size (386; 193 cases and 193 controls) was selected. Recently, the five referral hospitals found in Amhara national regional state are evenly distributed. They have their own NICU. Among the total of 2091 predicted number of newborns admitted to NICU irrespective of status of birth asphyxia, 193 newborns with birth asphyxia (cases) and 193 newborns without birth asphyxia (controls) were selected using systematic random sampling technique with proportional allocation. Every 2^nd^ of cases and every 8^th^ of controls were included in the study.

### Operational definition

Birth Asphyxia: it is the failure to initiate and sustain breathing at birth. Asphyxiated newborn not able to breath after birth and either convulsions/spasms or not able to suckle normally after birth or not able to cry after birth or Appearance Pulse Grimace Activity Respiration (APGAR) Score of < 7 [18].

### Data collection procedures and data quality control

The interviewer administered questionnaire was used for data collection. Through trained data collectors, the questionnaire was pretested in 5% of clients for possible modifications prior to data collection. The trained supervisors and the principal investigator supervised the data collection process. The collected data were checked daily for consistency, completeness, clarity and accuracy throughout the data collection process.

### Data analysis

Collected data were edited, coded and entered to Epi info version 7 software packages. These were then exported to Statistical Package for Social Sciences (SPSS) version 20 for analysis. First descriptive analysis was presented using frequency tables, figures, and percentages. In the second stage, by using logistic regression, bivariate logistic regression was fitted to screen candidate variables with *p-value*< 0.2 for the final model. Hosmer and Lemshow goodness of fit test was performed. Finally, multivariable logistic regression model through backward stepwise method was fitted to identify significant determinants of birth asphyxia. Adjusted Odds Ratio with 95% CI and *p-value*<0.05 were calculated to identify determinants of birth asphyxia among newborns.

## Results

### Socio-demographic and behavioral characteristics

A total of 193 asphyxiated newborn-mother pairs (cases) and 193 non-asphyxiated newborn-mother pairs (controls) were included in the study. Infants’ mothers mean (±SD) age was 26.63(±5.09) years. Among the total study groups, 109(56%) of cases and 105(54%) of controls were in the age group of 25-34 years. Majority of respondents (182 (94%) of cases and 170(88%) of controls) were married. More than half of the respondents; i.e. 103(53%) of cases and 121(63%) of controls were came from urban area. Housewives and farmers constituted 133(69%) of cases and 101(52%) of controls. Sixty three (33%) of cases and 53(28%) of controls were at elementary school (1-8 grades). One hundred thirteen (59%) of cases and 95 (49%) of controls had less than 2000 ETB monthly incomes. Most of the cases and controls were come to hospitals from the nearby areas. All respondents did not have smoking behavior. Majority of cases and controls (374 (97%)) hadn’t ever khat chewing behavior. However, 62 (32%) of cases and 65 (34%) of controls had ever drunk alcohol during the last pregnancy (Table 1).

**Table 1:** Socio-demographic and behavioral characteristics of respondents among newborns of Amhara Region Referral Hospitals, Ethiopia, 2018

| Variables | Controls (%)<br>(n=193) | Cases (%)<br>(n=193) | Total count (%)<br>(n=386) | Chi-square (p-value) |
| --- | --- | --- | --- | --- |
| Maternal age (in years) |  |  |  |  |
| 15-24 | 67 (34.7) | 63 (32.6) | 130 (33.7) | 0.198(0.906) |
| 25-34 | 105 (54.4) | 109 (56.5) | 214 (55.4) |  |
| 35-49 | 21 (10.9) | 21 (10.9) | 42 (10.9) |  |
| Marital status |  |  |  |  |
| Unmarried | 23 (11.9) | 11(5.7) | 34 (8.8) | 4.644(0.031) |
| Married | 170 (88.1) | 182 (94.3) | 352 (91.2) |  |
| Place of residence |  |  |  |  |
| Urban | 121 (62.7) | 103 (53.4) | 224 (58) | 3.446(0.063) |
| Rural | 72 (37.3) | 90 (46.6) | 162 (42) |  |
| Maternal occupation |  |  |  |  |
| Laborer & student | 28 (14.5) | 15 (7.8) | 43 (11.1) | 12.261(0.016) |
| Farmer | 33 (17.1) | 47 (24.4) | 80 (20.7) |  |
| Merchant | 27 (14.0) | 22 (11.4) | 49 (12.7) |  |
| Housewife | 68 (35.2) | 86 (44.6) | 154 (39.9) |  |
| Government employed | 37 (19.2) | 23 (11.9) | 60 (15.5) |  |
| Educational status |  |  |  |  |
| Unable to read and write | 39 (20.2) | 49 (25.4) | 88 (22.8) | 9.694(0.046) |
| Able to read and write with informal education** | 12 (6.2) | 17 (8.8) | 29 (7.5) |  |
| Elementary school (1-8 grades) | 53 (27.5) | 63 (32.6) | 116 (30.1) |  |
| High school and Prep(9-12 grades) | 42 (21.8) | 39 (20.2) | 81 (21.0) |  |
| Above 12 grades | 47 (24.4) | 25 (13.0) | 72 (18.7) |  |
| Monthly income (average in ETB) |  |  |  |  |
| <=2000 | 95 (49.2) | 113 (58.5) | 208 (53.9) | 3.384(0.184) |
| 2001-5000 | 73 (37.8) | 60 (31.1) | 133 (34.5) |  |
| >5000 | 25 (13.0) | 20 (10.4) | 45 (11.7) |  |
| Distance from the hospital (in KM) |  |  |  |  |
| 0-10 | 108 (56.0) | 78 (40.4) | 186 (48.2) | 12.441(0.002) |
| 11-50 | 62 (32.1) | 70 (36.3) | 132 (34.2) |  |
| 51-275 | 23 (11.9) | 45 (23.3) | 68 (17.6) |  |
| Khat chewing |  |  |  |  |
| No | 187 (96.9) | 187 (96.9) | 374 (96.9) | -* |
| Yes | 6 (3.1) | 6 (3.1) | 12 (3.1) |  |
| Alcohol drinking |  |  |  |  |
| No | 128 (66.3) | 131 (67.9) | 259 (67.1) | 0.106(0.745) |
| Yes | 65 (33.7) | 62 (32.1) | 127 (32.9) |  |

-\*chi-square assumption not fulfilled
- \*\*informal education includes education delivered with campaign and at religious institutes

### Maternal health related variables

Majorly 173 (90%) of cases and 177 (92%) of controls didn’t faced pre-eclampsia /eclampsia, however, 20 (10%) of cases and 16 (8%) of controls faced the problem. Eight (4%) of cases and 9(5%) of controls have HIV. Twenty two (9%) of cases and 14 (7%) of controls faced bleeding during their last pregnancy. Forty four (23%) of cases and 32(17%) of controls had iron-deficiency anemia. One hundred twenty four (64%) of cases and 89 (46%) of controls were referred from another health facilities (Table 2).

**Table 2:** Maternal health related variables among newborns of Amhara Region Referral Hospitals, Ethiopia, 2018

| Variables | Controls (%)<br>(n=193) | Cases (%)<br>(n=193) | Total count<br>(%)<br>(n=386) | Chi-square (p-value) |
| --- | --- | --- | --- | --- |
| Pre-eclampsia/eclampsia |  |  |  |  |
| No | 177 (91.7) | 173 (89.6) | 350 (90.7) | 0.490(0.484) |
| Yes | 16 (8.3) | 20 (10.4) | 36 (9.3) |  |
| HIV status |  |  |  |  |
| No | 184 (95.3) | 185 (95.9) | 369 (95.6) | 0.062(0.804) |
| Yes | 9 (4.7) | 8 (4.1) | 17 (4.4) |  |
| Diabetes Mellitus |  |  |  |  |
| No | 192 (99.5) | 191 (99.0) | 383 (99.2) | -* |
| Yes | 1 (0.5) | 2 (1.0) | 3 (0.8) |  |
| Bleeding in pregnancy (APH) |  |  |  |  |
| No | 179 (92.7) | 171 (88.6) | 350 (90.7) | 1.961(0.161) |
| Yes | 14 (7.3) | 22 (9.3) | 36 (9.3) |  |
| Iron-deficiency anemia |  |  |  |  |
| No | 161 (83.4) | 149 (77.2) | 310 (80.3) | 2.359(0.125) |
| Yes | 32 (16.6) | 44 (22.8) | 76 (19.7) |  |
| Referral status |  |  |  |  |
| No | 104 (53.9) | 69 (35.8) | 173 (44.8) | 12.832(0.000) |
| Yes | 89 (46.1) | 124 (64.2) | 213 (55.2) |  |

-\*chi-square assumption not fulfilled

### Antepartum and Intra-partum related variables

Among the total respondents, 91(47%) of mothers with cases and 105(54%) of mothers with controls experienced more than one pregnancies, however, 102(53%) of cases and 88(46%) of controls experienced their first pregnancies. Of the study units, 84(44%) of cases and 96(50%) of controls; and 109(57%) of cases and 97(50%) of controls were multiparous and primiparous respectively. Five (2.6%) of the cases and 16(8.3%) of controls had given twins during their last pregnancy. Nearly 189(98%) of cases and 188(97%) of controls had antenatal care (ANC) visits. Of the total respondents, 88(46%) of cases and 67(35%) of controls experience prolonged labor during the last pregnancy. Of the infants’ mothers, 45 (23%) of cases and 27 (14%) of controls faced premature rapture of membranes before labor starts. Only 17 (9%) of cases and 12 (6%) of controls faced prolonged rapture of membranes after 24 hours. Very few number of study subjects (11 (6%) of cases and 4 (2%) of controls) faced cord prolapse. Twelve (13%) of cases and 7 (4%) of controls had breech presentation. Of the total subjects, 53(28%) of cases and 61(32%) of controls were delivered with cesarean section, and 38(20%) of cases and 15(8%) of controls were delivered with instrumental assisted. Few respondents (30 (16%) of cases and 5 (3%) of controls) got the delivery service at health centers (Table 3).

**Table 3:** Antepartum and Intra-partum characteristics of respondents among newborns of Amhara Region Referral Hospitals, Ethiopia, 2018

| Variables | Controls (%)<br>(n=193) | Cases (%)<br>(n=193) | Total count (%)<br>(n=386) | Chi-square (p-value) |
| --- | --- | --- | --- | --- |
| Gravidity |  |  |  |  |
| Primigravida | 88 (45.6) | 102 (52.8) | 190 (49.2) | 2.032(0.154) |
| Multigravida | 105 (54.4) | 91 (47.2) | 196 (50.8) |  |
| Parity |  |  |  |  |
| Primiparous | 97 (50.3) | 109 (56.5) | 206 (53.4) | 1.499(0.221) |
| Multiparous | 96 (49.7) | 84 (43.5) | 180 (46.6) |  |
| Fetal outcome |  |  |  |  |
| Single | 188 (97.4) | 177 (91.7) | 365 (94.6) | 6.093(0.014) |
| Twins | 5 (2.6) | 16 (8.3) | 21 (5.4) |  |
| Number of ANC visits |  |  |  |  |
| 0 | 5 (2.6) | 4 (2.1) | 9 (2.3) | 7.579(0.023) |
| 1-3 | 78 (40.4) | 105 (54.4) | 183 (47.4) |  |
| >=4 | 110 (57.0) | 84 (43.5) | 194 (50.3) |  |
| Place of ANC visits |  |  |  |  |
| Private and Non-Governmental Organization's health facility | 28 (14.9) | 29 (15.3) | 57 (15.1) | 0.129(0.938) |
| Public health facility | 160 (85.1) | 160 (84.7) | 320 (84.9) |  |
| Prolonged labor |  |  |  |  |
| No | 126 (65.3) | 105 (54.4) | 231 (59.8) | 4.754(0.029) |
| Yes | 67 (34.7) | 88 (45.5) | 145 (40.2) |  |
| Premature rupture of membranes |  |  |  |  |
| No | 166 (86.0) | 148 (76.7) | 314 (81.3) | 5.532(0.019) |
| Yes | 27 (14.0) | 45 (23.3) | 72 (18.7) |  |
| Prolonged rupture of membranes |  |  |  |  |
| No | 181 (93.8) | 176 (91.2) | 357 (92.5) | 0.932(0.334) |
| Yes | 12 (6.2) | 17 (8.8) | 29 (7.5) |  |
| Cord prolapse |  |  |  |  |
| No | 189 (97.9) | 182 (94.3) | 371 (96.1) | 3.399(0.065) |
| Yes | 4 (2.1) | 11 (5.7) | 15 (3.9) |  |
| Presentation |  |  |  |  |
| Cephalic | 186 (94.4) | 168 (87.0) | 354 (91.7) | 11.040(0.001) |
| Breech | 7 (3.6) | 25 (13.0) | 32 (8.3) |  |
| Mode of delivery |  |  |  |  |
| Vaginally | 117 (60.6) | 102 (52.8) | 219 (56.7) | 11.570(0.003) |
| Cesarean section | 61 (31.6) | 53 (27.5) | 114 (29.5) |  |
| Instrumental | 15 (7.8) | 38 (19.7) | 53 (13.7) |  |
| Place of delivery |  |  |  |  |
| Health center | 5 (2.6) | 30 (15.5) | 35 (9.1) | 19.638(0.000) |
| Hospital | 188 (97.4) | 163 (84.5) | 351 (90.9) |  |

### Newborn characteristics

Of the total newborns, 117(61%) of cases and 104(54%) of controls are males. Among all sexes, 65(35%) of cases and 10(5%) of controls had low birth weight. Both preterm and post-terms contributed 43(22%) of cases and 17(9%) of controls. One hundred forty six (76%) of cases were unable to breath after birth, however, only 31(16%) of cases experienced spasm. The majority, 189 (98%) of cases were unable to suckle normally after birth, and 184 (95%) of cases were unable to cry after birth (Table 4).

**Table 4:** Newborn characteristics among newborns of Amhara Region Referral Hospitals, Ethiopia, 2018

| Variables | Controls (%)<br>(n=193) | Cases (%)<br>(n=193) | Total count (%)<br>(n=386) | Chi-square (p-value) |
| --- | --- | --- | --- | --- |
| Sex of newborn |  |  |  |  |
| Male | 104 (53.9) | 117 (60.6) | 221 (57.3) | 1.789(0.181) |
| Female | 89 (46.1) | 76 (39.4) | 165 (42.7) |  |
| Birth weight |  |  |  |  |
| Low birth weight (<2.5kg) | 10 (5.2) | 67 (34.7) | 77 (19.9) | 52.709(0.000) |
| Normal (≥2.5kg) | 183 (94.8) | 126 (65.3) | 309 (80.1) |  |
| Gestational age (Wks) |  |  |  |  |
| <37 (Preterm) | 5 (2.6) | 32 (16.6) | 37 (9.6) | 21.820(0.000) |
| 37-41(Term) | 176 (91.2) | 150 (77.7) | 326 (84.5) |  |
| >41 (Post-term) | 12 (6.2) | 11 (5.7) | 23 (6.0) |  |
| Able to breath after birth |  |  |  |  |
| No | 0 (0.0) | 146 (75.6) | 146 (37.8) | -* |
| Yes | 193 (100) | 47 (24.4) | 240 (62.2) |  |
| Had convulsions / spasm |  |  |  |  |
| No | 193 (100) | 162 (83.9) | 355 (92.0) | -* |
| Yes | 0 (0.0) | 31 (16.1) | 31 (8.0) |  |
| Able to suckle normally after birth |  |  |  |  |
| No | 0 (0.0) | 189 (97.9) | 189 (49.0) | -* |
| Yes | 193 (100) | 4 (2.1) | 197 (51.0) |  |
| Able to cry after birth |  |  |  |  |
| No | 0 (0.0) | 184 (95.3) | 184 (47.7) | -* |
| Yes | 193 (100) | 9 (4.7) | 202 (52.3) |  |
| APGAR Score |  |  |  |  |
| Severe (0-3) | 0 (0.0) | 13 (6.7) | 13 (3.4) | -* |
| Mild to moderate (4-6) | 0 (0.0) | 180 (93.3) | 180 (46.6) |  |
| Normal (7-10) | 193 (100) | 0 (0.0) | 193 (50.0) |  |

-\*chi-square assumption not fulfilled

### Determinants of birth asphyxia

In binary logistic regression analysis, twenty seven variables were entered in the analysis and only twenty variables were identified as determinants of birth asphyxia (Table 6). The others theme of variables did not have association to birth asphyxia. Variables that have p< 0.2 in the bivariate analysis and enter to multivariable analysis were; maternal marital status, place of residence, occupation, education, distance from the hospitals, bleeding during pregnancy, iron-deficiency anemia, referral status, gravidity, multiple births, number of ANC visits, prolonged labor, premature rapture of membrane, cord prolapse, fetal presentation, mode of delivery, place of delivery, gestational age, sex of newborn, and birth weight. After adjustment, the determinants that have p<0.05 at 95% confidence interval are only prolonged labor, mode of delivery, place of delivery, and birth weight (Table 5).

**Table 5:** Determinants of birth asphyxia among newborns of Amhara Region Referral Hospitals, Ethiopia, 2018

| Variables | Controls | Cases | Crudes and Adjusted Odds Ratios with 95% Confidence Intervals |  |
| --- | --- | --- | --- | --- |
|  |  |  | COR (95% CI) | AOR (95% CI) |
| Marital status |  |  |  |  |
| Unmarried | 23 | 11 | 1 | 1 |
| Married | 170 | 182 | 2.24(1.06, 4.73) | 1.79(0.71, 4.47) |
| Place of residence |  |  |  |  |
| Urban | 121 | 103 | 1 | 1 |
| Rural | 72 | 90 | 1.47(0.98, 2.21) | 0.78(0.36, 1.66) |
| Maternal occupation |  |  |  |  |
| Laborer & student | 28 | 15 | 0.86(0.38, 1.95) | 1.04(0.39, 2.83) |
| Farmer | 33 | 47 | 2.29(1.16, 4.54)* | 1.74(0.67, 4.49) |
| Merchant | 27 | 22 | 1.31(0.61, 2.82) | 0.90(0.33, 2.42) |
| Housewife | 68 | 86 | 2.04(1.11, 3.74)* | 1.52(0.70, 3.30) |
| Government employed | 37 | 23 | 1 | 1 |
| Educational status |  |  |  |  |
| Unable to read and write | 39 | 49 | 2.36(1.24, 4.49)* | 1.83(0.51, 6.59) |
| Able to read and write | 12 | 17 | 2.66(1.10, 6.45)* | 2.07(0.51, 8.42) |
| Elementary school (1-8 grades) | 53 | 63 | 2.24(1.22, 4.10)* | 1.73(0.57, 5.27) |
| High school and Prep(9-12 grades) | 42 | 39 | 1.75(0.91, 3.35) | 1.78(0.62, 5.14) |
| Above 12 grades | 47 | 25 | 1 | 1 |
| Distance from the hospital (in KM) |  |  |  |  |
| 0-10 | 108 | 78 | 1 | 1 |
| 11-50 | 62 | 20 | 1.56(0.99, 2.45) | 0.74(0.39, 1.39) |
| 51-275 | 23 | 45 | 2.71(1.52, 4.84)* | 1.69(0.78, 3.65) |
| Bleeding in pregnancy (APH) |  |  |  |  |
| No | 179 | 171 | 1 | 1 |
| Yes | 14 | 22 | 1.65(0.82, 3.32) | 1.24(0.51, 3.01) |
| Iron-deficiency anemia |  |  |  |  |
| No | 161 | 149 | 1 | 1 |
| Yes | 32 | 44 | 1.49(0.90, 2.47) | 1.29(0.68, 2.44) |
| Referral status |  |  |  |  |
| No | 104 | 69 | 1 | 1 |
| Yes | 89 | 124 | 2.10(1.40, 3.16)* | 1.72(0.95, 3.11) |
| Gravidity |  |  |  |  |
| Primigravida | 88 | 102 | 1 | 1 |
| Multigravida | 105 | 91 | 0.75(0.50, 1.12) | 0.83(0.48, 1.42) |
| Fetal outcome |  |  |  |  |
| Single | 188 | 177 | 1 | 1 |
| Twins | 5 | 16 | 3.40(1.22, 9.47)* | 1.90(0.51, 7.15) |
| Number of ANC visits |  |  |  |  |
| 0 | 5 | 4 | 1.05(0.27, 4.02) | 0.22(0.04, 1.15) |
| 1-3 | 78 | 105 | 1.76(1.17, 2.65)* | 1.34(0.81, 2.21) |
| >=4 | 110 | 84 | 1 | 1 |
| Prolonged labor |  |  |  |  |
| No | 126 | 105 | 1 | 1 |
| Yes | 67 | 88 | 1.58(1.05, 2.38)* | 2.00(1.20, 3.36)** |
| Premature rupture of membranes |  |  |  |  |
| No | 166 | 148 | 1 | 1 |
| Yes | 27 | 45 | 1.87(1.11, 3.16)* | 1.55(0.82, 2.95) |
| Cord prolapse |  |  |  |  |
| No | 189 | 182 | 1 | 1 |
| Yes | 4 | 11 | 2.86(0.89, 9.13) | 3.41(0.86, 13.46) |
| Presentation |  |  |  |  |
| Cephalic | 186 | 168 | 1 | 1 |
| Breech | 7 | 25 | 3.95(1.67, 9.38)* | 2.71(0.96, 7.70) |
| Mode of delivery |  |  |  |  |
| Spontaneous vaginal delivery (SVD) | 117 | 102 | 1 | 1 |
| Cesarean section | 61 | 53 | 0.99(0.63, 1.57) | 0.96(0.55, 1.67) |
| Instrumental | 15 | 38 | 2.91(1.51, 5.59)* | 3.03(1.41, 6.49)** |
| Place of delivery |  |  |  |  |
| Health center | 5 | 30 | 6.92(2.62, 18.25)* | 7.36(2.44, 22.13)*** |
| Hospital | 188 | 163 | 1 | 1 |
| Sex of newborn |  |  |  |  |
| Male | 104 | 117 | 1 | 1 |
| Female | 89 | 76 | 0.76(0.51, 1.14) | 0.79(0.48, 1.30) |
| Birth weight |  |  |  |  |
| Low birth weight (<2.5kg) | 10 | 67 | 9.73(4.82, 19.64)* | 8.94(4.08, 19.56)*** |
| Normal (≥2.5kg) | 183 | 126 | 1 | 1 |
| Gestational age (Wks) |  |  |  |  |
| <37 (Preterm) | 5 | 32 | 6.98(2.00, 24.32)* | 4.02(0.89, 18.13) |
| 37-41(Term) | 176 | 150 | 0.93(0.40, 2.17) | 0.70(0.26, 1.93) |
| >41 (Post-term) | 12 | 11 | 1 | 1 |

\* p<0.20; \*\*p<0.01; \*\*\*p<0.001

Newborns born from mothers with prolonged labor were 2 times more likely to suffer from birth asphyxia as compared to their counterparts (AOR: 2.00, 95% CI: 1.20, 3.36). Newborns that were born using instrumental delivery were 3.03 times more likely to develop birth asphyxia than those delivered by vaginally (AOR: 3.03, 95% CI: 1.41, 6.49). Newborns that were born at health centers were 7.36 times more likely to develop birth asphyxia than those born at hospitals (AOR: 7.36, 95% CI: 2.44, 22.13). Newborns with low birth weight (2.5kg) had 8.94 higher odds of birth asphyxia than those of normal (>=2.5kg) at birth (AOR: 8.94, 95% CI: 4.08, 19.56) (Table 5).

## Discussion

This study identified significant determinants of birth asphyxia in Amhara National Regional State Referral Hospitals, Ethiopia. Some of intra-partum and newborn-related variables were associated to birth asphyxia. Prolonged labor, mode of delivery (instrumental), place of delivery (at health centers), and low birth weight were identified as significant determinants of neonatal birth asphyxia. This study will be informing health care providers especially at health centers for their appropriate interventions and even it may needs managerial decisions for improving referral systems to make fast to reduce the burden of birth asphyxia occurred at primary health care level. This finding shows that prolonged labor is statistically significant determinant of birth asphyxia. It is in line with a cross-sectional study done in Jimma zone of Ethiopia [19]. Other previous studies have also shown similar results [20, 21]. Women with a prolonged labor had a negative birth experience more often than did women who had a normal labor [22]. According to American Pregnancy Association and Reiter and Walsh, PC, prolonged labor or failure to progress occurs when labor lasts for approximately 20 hours or more if you are a first-time mother, and 14 hours or more if you have previously given birth. A prolonged latent phase happens during the first stage of labor. It can be exhausting and emotionally draining, but rarely leads to complications. Prolonged labor may happen due to slow effacement of the cervix, too large baby, too small birthing canal or woman’s pelvis, carrying multiples, incorrect fetal presentation, psychological factors, such as worry, stress, or fear [23, 24].

Our study result shows that mode of delivery (in our case, instrumental delivery) determined the occurrence of neonatal birth asphyxia. This in agreement with a case control study done in India [25] and cross-section findings in Ethiopia [26] and Pakistan [27]. A research conducted in England revealed that infants born by instrumental delivery (forceps and vacuum delivery) for presumed fetal compromise had the poorest condition at birth [28]. Infants delivered by instrumental delivery had the worst neonatal effects, suggesting the mode of delivery itself is influential [28]. Instrumental delivery is permitted when spontaneous vaginal delivery is failed. It is, therefore, the practitioners may delay to practice after the rapture of membrane and the newborn may come to asphyxia.

The study shows that neonates born at health centers had higher risk of birth asphyxia than those who born at hospitals. In many scholars, place of delivery, in general, has an association with birth asphyxia [14, 29, 30]. In our cases we couldn’t get similar studies for comparison, however, there might be different causes of higher asphyxiated cases born in health centers, such as lack of skilled birth attendants in the health centers, and/or delay to refer the cases to hospitals, and/or transportation issues.

This finding shows that neonates with low birth weight had higher risk of asphyxia than those with normal birth weight. Other studies in Thailand, Pakistan, and Iran also revealed similar results [10, 14, 17]. Low birth weight is mostly indicated as a fetal risk factor. The primary cause of low birth weight is premature birth (being born before 37 weeks gestation), as it is true for our finding, 10% of neonates were preterm. Another causes of low birth weight is intrauterine growth restriction, maternal health issues, early maternal age, and multiple births [31]. Low birth weight neonates should to be given much more attention compared to their counterparts whose birth weight are normal as they are prone to asphyxia [32].

This study has strength in that the study was done at region level on five referral hospitals, which may reflect regional burden at hospital levels. There are limitations of the study as it is hospital based study where majority of births were attended by qualified personnel, this does not reflect exact risk factors prevalent in the community, where majority of births are unable to access those referral hospitals. Also, because a person is assigning the number, the Apgar score is subjective that may under or overestimate the magnitude of birth asphyxia.

## Conclusions

This study identified determinants of neonatal birth asphyxia in Amhara National Regional State Referral Hospitals, Ethiopia. Prolonged labor/failure to progress, mode of delivery (instrumental), place of delivery (at health centers), and low birth weight were identified as statistically significant determinants of birth asphyxia. Even though most of the identified variables are the common and familiar causes of birth asphyxia, neonates born at health centers were more exposed to birth asphyxia than neonates born in hospitals. This might be due to delay of referral process and lack of skilled professionals in health centers. Consequently, it may indicate the need of operational intervention planning and further researches. Further researches may be recommended to identify why neonatal birth asphyxia is high at health centers than hospitals.

### Ethics approval and consent to participate

Ethical clearance was obtained from the ethical review board of School of Public Health, College of Medicine and Health Sciences, Bahir Dar University. Permission was obtained from Amhara public health institute and five referral hospitals. Full explanation of the study was given to respondents and oral consent was obtained from them. Personal identifiers (like respondents’ name) were not included in data collection. That is confidentiality of data was maintained anonymously. There was no known risk on participating in this study. Study subjects might not directly benefit from participating in the study. However, information obtained from this study may be used to improve the health of newborns. Respondents were free to decline participation or withdraw from study participation during data collection.

## Authors’ contributions

**AD** conceived the concept of the study, designed the study, and prepared the proposal, involved in the data analysis and interpretation. **MA** and **EW** involved in the designing of the study, revised the proposal, involved in the data analysis and interpretation. **GG** assisted and provided technical support on every step of proposal development and data management. All of the authors contributed to the preparation of the manuscript and approved the final version for publication.

## Supporting information

S1 Data. This is the data set of the study. (SAV)

## Acknowledgements

We are very grateful to Bahir Dar University for the approval of the ethical clearance. We would also like to thank all individuals participated in this study for their cooperation in taking part in this study.

